# Genomic feature of filled regions of telomere-to-telomere genomes in five model species

**DOI:** 10.1101/2024.12.31.630859

**Authors:** Chu Xiong, Jun Zheng, Hui Zhang, Yunpeng Zhang, Lihong Hao, Min Wang, Zhen Liang

**Affiliations:** College of Biomedical Engineering, Taiyuan University of Technology, Taiyuan 030024, China; Institute of Wheat Research, Shanxi Agricultural University, Linfen 041000, China; College of Life Sciences, Shanxi University, Taiyuan 030006, China

**Keywords:** Telomere to Telomere, filled regions, genomic feature, model species

## Abstract

T2T genome is a telomere-to-telomere high-quality genome with exceptional accuracy, continuity and integrity. It significantly filled some gaps of chromosome. However, the genomic features unique to the filled regions assembled in the T2T genome across species remain unclear. Here, we collected and compared T2T genomes with their older version from human (*Homo sapiens*), animal (*Mus musculus*), and plants (*Arabidopsis thaliana*, *Musa acuminata*, and *Fragaria vesca*). We explored the newly filled regions generated by the T2T genome in comparison to the old one, and focused on: (1) the comparison of their locations, (2) the distribution of genomic features, and (3) gene annotation and gene function for filled regions. Research indicated that the filled regions of the T2T genome ordinarily appear in the centromere and telomere regions, contain a great number of repetitive sequences. The T2T genome revealed the identification of a certain number of newly annotated genes, alongside significant variations in common annotated genes. Interestingly, the genomic features of filled regions exhibit distinct patterns between plants and human/animal. T2T genomes feature analysis serves as a crucial resource for studying pattern characteristics across diverse species and advancing the in-depth development of genomics.

## 1. Introduction

The genome encapsulates fundamental genetic information within an organism, including protein coding genes, non-coding gene, regulatory elements, and repetitive sequences within the DNA molecules and so on [1]. Genome assembly and annotation play crucial roles in exploring genome and transcriptome comparisons [2], gene expression [3, 4] and species evolution [5]. Reference genome assembly from sequence data facilitates the identification of different expression patterns from annotated genes, as opposed to de novo assembly for transcripts [2]. In evolution research, genomic analysis aids in determining genetic relatedness and identifying genomic features conducive to computational design of artificial overlapping genes [6, 7]. The assembled genome sequence plays important roles in understanding biological function, disease pathogenesis and agricultural improvement.

The advent of high-throughput sequencing technologies has significantly advanced whole genome sequencing and complete genome assembly [8]. Currently, short-and long-read sequencing data from high-throughput sequencing platforms are extensively employed to study genome structure, function, evolution, gene mapping, and genetic polymorphisms [9]. Notably, complete genome assembly involves assembling fragments contigs from raw whole-genome sequencing data into a coherent complete genomic sequence [10–12]. The complete genome assembly offers multiple advantages, including obtaining contiguous and complete genome sequences, discovering new genes and functions, elucidating genome structural variations and evolutionary history, and facilitating genome annotation [13, 14].

Previous studies have extensively explored features of genomic sequences, such as GC content [15], encoded protein sequence [16], scaffold N50 length [17] and identification of repeats [18, 19]. Regions with high GC content in gene tend to exhibit higher gene densities and longer open reading frames (ORFs) [20]. Mutations and polymorphisms in coding protein sequences are often associated with diseases, such as cancer and genetic disorders [16], impacting protein structure, function, and interactions [21]. Scaffold N50 is a crucial metric for assessing the quality of genome assembly, with higher values indicating improved continuity and completeness of the assembled genome [17]. Additionally, repetitive sequences in the genome play significant roles in species evolution, leading to highly conserved or rapidly evolving regions [18, 19]. In depth analysis of sequence features is closely linked to gene function, with biological function annotation involving the identification of regulatory elements, including promoter and enhancer regions [22, 23], and non-coding genes within the genome sequence [24]. The relationship between germline genes and human cancer is intricately connect to gene family expansion and evolutionary process [25]. Analyzing and summarizing of sequence features provided insights into genome characteristics and functions, aids in locating and annotating of unknown genes, and enhances the understanding of biology and molecular evolution.

To date, several telomere to telomere (T2T) complete genomes have been assembled and are available in public databases, including *Homo sapiens* [26, 27], *Mus musculus* [28, 29], *Musa acuminata* [30, 31], *Arabidopsis thaliana* [32, 33] and *Fragaria vesca* [34, 35]. T2T genomes have significantly improved the DNA sequence integrity and exhibit higher functional diversity compared to previous genome versions with gaps. For instance, the release of the human T2T-CHM13 [26] genome has spurred a series of comparative studies between T2T and earlier genome assemblies. In comparison to GRCh38 [27], T2T-CHM13 has identified over 1 million whole-genome variants [36]. This discovery holds significant implications for the comprehensive detection of fragment repeats, intricate compound repeat structures, comprehensive gene identification, and the elucidation of genetic variation patterns [37–39]. Moreover, T2T-CHM13 includes a complete paracentric centromere DNA sequence, facilitating detailed computational and analytical studies of human centromere organization and genetic information [40].

However, the similarities and differences of genomic features in the filled regions (FR) that unique assembled in T2T genome across different species remain unclear. This study comprehensively investigates the specific properties of T2T genome across different species, analyzing and comparing human, animal and plant T2T genomes in four aspects: basic sequence information, distribution of FR, characteristic analysis and gene function. This research holds profound significance for advancing genomic studies and exploring evolutionary patterns.

## 2. Materials and methods

### 2.1 Data collection

The latest complete T2T genomes from human, animal and plant species were obtained from various sources: NCBI (*H. sapiens*), GENCODE (*M. musculus*), GitHub (*A. thaliana*) and respective genome websites (*M. acuminata* and *F. vesca*) (Table 1). The previous version of these five genomes, which included gaps, were acquired from NCBI (*H. sapiens, A. thaliana, M. acuminata*), GENCODE (*M. musculus*) and genome websites (*F. vesca*) and are referred to as ‘old’ genomes. The human T2T genome CHM13v2.0, consisting of 22 autosomes, X and Y chromosome assembled by human T2T Consortium [26], was analyzed and compared with previous old genome GRCh38.p14 [27]. The gapless version of mouse genome (GRCm39) [41] was compared with the previous iteration (GRCm38.p6). In the case of plants, the *A. thaliana* genomes TAIR10.1 [32] and Col-CEN_v1.2 [33], *M. acuminata* genomes ASM31385v2 [30] and Musa_pahang_v4 [31] and *F. vesca* genomes F._vesca_v4 [34] and F._vesca_v6 [35] were collected and subjected to comparative analysis (Table 1). In addition, we compared the basic statistical metrics, including genome size, GC content and scafford N50 between all T2T genomes and old genomes.

**Table 1.**
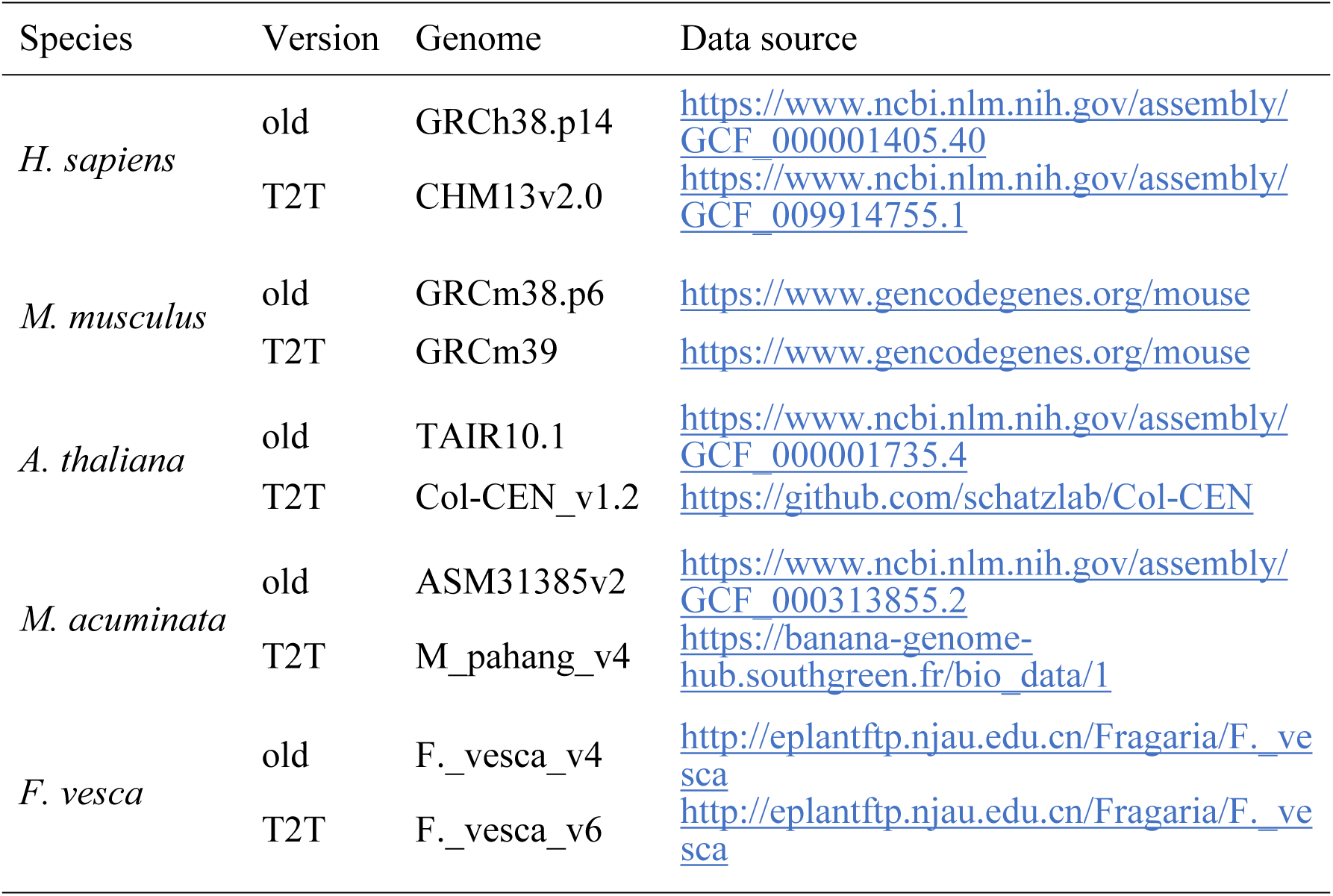
T2T and old genomes of five species.

### 2.2 Identification of FR in five T2T genomes

The FR refers to the uniquely assembled sequences in T2T genome compared to the old genome in five species. Regions where the old genome aligns to the T2T genome are considered shared regions, while the remaining regions in the T2T genome, relative to the old genome as FR. Sequence comparison between T2T and old genomes was performed using the ‘nucmer’ of MUMmer (http://mummer.sourceforge.net) [42]. The T2T genome serve as the reference genome, and the old genome was as the query sequence aligned against the reference genome. Low-quality alignment results were filtered with ‘-l’ parameters. FR in 23 chromosomes (22 autosomes and X) of *H. sapiens*, 21 chromosomes (19 autosomes, X and Y) of *M. musculus*, 5 chromosomes of *A. thaliana*, 12 chromosomes (11 chromosomes and mitochondria) of *M. acuminata* and 7 chromosomes of *F. vesca* were identified and compared, respectively. Genomic features within FR were analyzed and compared between T2T and old genomes, encompassing location, sequences features and annotation genes within the FR. The results of the global collinearity analysis for five species were visualized through mummerplot completion.

### 2.3 Comparison of FR location

The distribution patterns of FR resulting from the alignment of T2T and old genomes was performed and compared in five species. The proportional representation of FR in each genome was calculated by finding the complement region of the aligned regions on the chromosomes using BEDtools (https://bedtools.readthedocs.io) [43]. Utilizing the positional information of FR and aligned regions, we acquired global locus information for each chromosome in each T2T genome by merging these two regions. The plots illustrating the distribution of FR in each genome were generated using ggplot2 package [44] of R language v.4.3.1 [45].

For *H. sapiens*, *A. thaliana* and *F. vesca* genomes, the start and stop sites of centromeres were determined from their T2T genomes’ annotation files. Additionally, we identified the start and stop sites of telomeres for each chromosome in *F. vesca* using its T2T genomic information. Combining this information with the location of FR to calculate the proportion of FR within all known centromere and telomere regions. Based on the genome annotation files of the T2T genomes [26, 31, 33, 35, 41], we analyzed the number and proportion of gene regions and intergenic regions in the FR, respectively. For gene regions, the location information of CDS, exon and intron was obtained after analysis using GenomicFeatures package [46] in R language [45]. Regarding intergenic regions, all location information of ‘gene’annotation type was extracted from genome annotation files, and the complementary regions were then acquired as the location information of ‘intergenic’ regions. Finally, the number and proportion of each type in the FR were calculated using BEDtools [43].

### 2.4 Comparison of filled sequences features

To delve deeper into the characteristics of FR sequences, we annotated DNA repetitive sequences from T2T genomes using RepeatMasker (https://www.repeatmasker.org) [47]. For *H. sapiens*, *M. musculus*, and *A. thaliana*, we utilized a combination of the RepBase (v.20181026) [48] and Dfam (v.20170127) [49] repeat databases for dentifying and annotating repeat sequences. Due to the absence of candidate repeat databases for *M. acuminata* and *F. vesca*, RepeatModeler (v.2.0.1) [50] was utilized to identify and model repetitive elements or sequences in their genomes. Subsequently, RepeatMasker was performed with the Dfam (v.20230110) database configured to identify and annotate repeat sequences. The proportion of repeat sequences within the FR on each chromosome was calculated using BEDTools [43]. Furthermore, we analyzed the types of repeat sequences in the T2T genomes and their positional distribution throughout the entire genome, and compared them with the distribution of the FR. The proportion and position distribution of different types of repeat types within the FR of each chromosome were visualized using ggplot2 [44] in the R language.

### 2.5 Gene annotation of filled sequences

Functional analysis of FR from T2T genomes is crucial for understanding assembly improvement and biological research. Genes are the most direct and important entities for functional analysis of FR. Information on identified genes, CDS and other annotations within the FR was obtained through the annotation files of T2T genomes [26, 31, 33, 35, 41] using BEDTools [43]. To calculate and analyze newly annotated genes in T2T genomes, we compared gene annotations from the annotation files of the T2T genomes [26, 31, 33, 35, 41] with those of old genomes in *H. sapiens*, *M. musculus* and *A. thaliana*. We obtained common annotated genes and newly annotated genes in the T2T genomes. We analyzed the number of extended genes and shortened genes, maximum extension and shortening length, and other information in the T2T genome. For the newly annotated genes, we calculated the number and proportion of repeated sequences to analyze the sequence characteristics of the newly annotated gene regions using BEDTools [43]. KEGG enrichment analysis was performed on the newly annotated genes of *M. musculus* using cluster Profiler [51] package in the R language to identify specific function in these newly annotated genes.

To further validate gene function and explore additional characteristics within the FR, we collected RNA-seq datasets corresponding to the five species and align them to T2T genomes using bwa v.0.7.17 [52] (Table S1). The mapping ratio (MP) of RNA-seq datasets on the T2T genome were calculated across five species. Additionally, we conducted a thorough analysis of the number of aligned reads from RNA-seq data within these FR. The alignments of RNA-seq datasets enhance the discovery potential of new transcripts and regulatory elements that may exist within the FR.

### 2.6 Gene function prediction and reliable FR extraction for *H. sapiens*

For *H. sapiens*, in addition to comparing the T2T and old genome versions using MUMmer [42], we also performed a comparison with minimap2 [53]. The new regions identified by each alignment were labeled as FR_1 and FR_2, respectively, and the overlapping regions between these were extracted as the final result FR. We utilized the resulting FR for motif analysis using Homer (http://homer.ucsd.edu) with the ‘findMotifsGenome.pl’ command. This command identifies and analyzes transcription factor binding sites and other functional elements across the entire genome. Subsequently, we confirmed potential gene functions within FR. In the ‘knownResults.txt’ file, we quantified the number of motifs with significant p-values and conducted KEGG enrichment pathway analysis on the genes associated with these significant motifs by clusterProfiler [51] package in the R language. This analysis provided a clearer understanding of how these genes are involved in specific biological pathways. Additionally, we extracted key pathways from the enrichment analysis results to elucidate the roles of these genes in cellular functions and disease mechanisms. Furthermore, we extracted sequences aligned to FR from all RNA-seq samples in 2.5 for the purpose of obtaining reliable FR sequences. We then aligned these sequences to the Nucleotide Collection (NT) database [54] using ‘blastn’ in BLAST v2.11.0 (https://blast.ncbi.nlm.nih.gov/Blast.cgi). This allowed us to identify homologous sequences of FR in other genomes and species. We extracted sequences aligned to non-*H. sapiens* species, realigned these sequences to the prior GRCh38 version, and selected unaligned sequences for use as reliable FR sequences. These sequences underwent Sanger sequencing [55] to verify their authenticity and accuracy.

### 2.7 Sequencing validation of reliable FR sequences of the T2T genome for *H. sapiens*

Through the reliable extraction of FR sequences in step 2.6, three sequences were ultimately selected for Sanger sequencing. To enhance the accuracy and coverage of sequencing, while avoiding sequence loss due to read length limitations or edge effects, and to provide more comprehensive upstream and downstream sequence information, approximately 300 bp were extended at both the 5’ and 3’ ends of each sequence. Subsequently, forward and reverse primers were designed for each sequence, and HeLa and HEK293T cell lines of H. sapiens were selected as experimental models for PCR (Polymerase Chain Reaction) [56] amplification, product detection, and purification. Primers were designed by the software Primer Premier Version 5.0 (Premier Biosoft International, Palo Alto, CA, USA), and the primers were synthesized by Sangon (https://www.sangon.com). The Primer sequences are shown in Table S2. PCR were performed in total volumes of 15 μl, including 3 pmol of each primer, 120 μM of each dNTP, 80ng of template DNA or cDNA, 0.75U of La-Taq, and 7.5 μl of 2 × buffer [TaKaRa Biotechnology (Dalian) Co. Ltd]. PCRs were performed as follows: 95 °C for 4 min; followed by 30–35 cycles of 95 °C for 30 s, annealing (55–62 °C) for 30 s, and extension at 72 °C (30 s to 3min), and 72 °C for 30 s, with a final extension of 72 ° C for 10min. The annealing temperatures and extension times depended on the primer sets and lengths of expected PCR products.

Finally, the PCR products were sent to Sangon Biotech Co., Ltd. (Shanghai) for Sanger bidirectional sequencing using the 3730XL sequencer. The sequencing process generated peak chromatograms in ‘.ab1’ format and sequence files in ‘.seq’ format. We conducted a thorough analysis of the electrophoretic bands produced during sequencing, as well as the final chromatograms, and validated the sequencing results to ensure accuracy and reliability.

## 3. Results

### 3.1 Overview of T2T genomes in five species

The new genomes of human, animal and plants species have seen increases compared to the old genomes. The genome size of *M. musculus*, *F. vesca*, *H. sapiens*, *A. thaliana*, *M. acuminata* in the T2T genomes exhibits increases of 0.09%, 0.14%, 0.72%, 10.92%, and 44.42%, respectively, when compared to the old genomes. Notably, *M. acuminata* demonstrates the largest increase in genome size. For GC content, there is a slight increase in *M. musculus*, *A. thaliana* and *F. vesca*. Regarding the assembly N50, the T2T genomes of all species exhibit improvements compared to the old genomes, which is closely related to the promotion of contig number in T2T genomes. The enhancement is particularly prominent in *M. acuminata*, which shows 24.6 times increase compared to the old genome (Table 2).

**Table 2.** Comparison of old and T2T genomes in five species.

| Species | Genome | Size (Mb) | GC (%) | Contig # | Assembly N50 (Mb) | Chr. # | Ratio of FR (%) |
| --- | --- | --- | --- | --- | --- | --- | --- |
| *H. sapiens* | Old_homo | 3,095.0 | 41.49 | 985 | 67.8 | 24 | 6.45 |
|  | T2T_homo | 3,117.3 | 40.74 | 24 | 150.6 | 24 |  |
| *M. musculus* | Old_mus | 2,647.5 | 41.50 | 139 | 54.5 | 21 | 4.35 |
|  | T2T_mus | 2,649.9 | 42.00 | 61 | 106.1 | 21 |  |
| *A. thaliana* | Old_arab | 119.1 | 36.05 | 7 | 23.5 | 5 | 10.10 |
|  | T2T_arab | 132.1 | 36.24 | 7 | 25.7 | 5 |  |
| *M. acuminata* | Old_musa | 331.8 | 40.73 | 7,512 | 1.3 | 11 | 26.38 |
|  | T2T_musa | 479.2 | 39.08 | 124 | 33.6 | 11 |  |
| *F. vesca* | Old_frag | 220.5 | 38.35 | 61 | 7.9 | 7 | 1.43 |
|  | T2T_frag | 220.8 | 38.50 | 7 | 34.34 | 7 |  |

For each species analyzed, T2T genomes typically exhibit higher ’C’ values, particularly noticeable in *M. acuminata*, and *F. vesca*. Moreover, T2T genomes tend to demonstrate lower ’F’ and ’M’ values in most instances, as observed in *M. musculus*, *M. acuminata*, and *F. vesca*. For both *H. sapiens* and *A. thaliana*, the BUSCO scores for the T2T and old genome versions were essentially identical. These findings suggest that T2T genomes generally possess superior genomic integrity. This enhancement in quality highlights the potential benefits of T2T methodologies in improving the continuity of genome assemblies (Table 3).

**Table 3.** Comparison of BUSCO scores between old and T2T genomes.

| Species | Genome | C | D | F | M | n |
| --- | --- | --- | --- | --- | --- | --- |
| *H. sapiens* | Old_homo | 99.2% | 0.7% | 0.2% | 0.7% | 13,780 |
|  | T2T_homo | 99.2% | 0.7% | 0.2% | 0.7% | 13,780 |
| *M. musculus* | Old_mus | 99.1% | 11.9% | 0.4% | 0.5% | 954 |
|  | T2T_mus | 99.1% | 11.5% | 0.4% | 0.5% | 954 |
| *A. thaliana* | Old_arab | 100% | 16.5% | 0.0% | 0.0% | 255 |
|  | T2T_arab | 100% | 16.5% | 0.0% | 0.0% | 255 |
| *M. acuminata* | Old_musa | 98.4% | 23.1% | 1.2% | 0.4% | 255 |
|  | T2T_musa | 98.8% | 24.7% | 0.8% | 0.4% | 255 |
| *F. vesca* | Old_frag | 98.7% | 1.9% | 0.5% | 0.8% | 1,614 |
|  | T2T_frag | 98.8% | 3.6% | 0.5% | 0.7% | 1,614 |

(C: complete; D: complete and duplicate; F: fragment; M: missing; n: total groups)

Comparing the T2T genomes to the old genomes, there is a general trend of larger genome sizes and higher assembly N50 values (Table 2). Additionally, the degree of differences between the T2T and old genomes varies among the five species, with distinct attributes showing improvement in the T2T genomes for each species. The differences between the T2T and old genomes in *M. musculus* are comparatively smaller than in other species, while in *M. acuminata*, they are more substantial. The assembly N50 of the *H. sapiens* has a significant increase, whereas the increase in *A. thaliana* is not as pronounced.

### 3.2 The distribution of FR

The proportion and the number of FR vary among the five T2T genomes. T2T_musa exhibits the highest proportion of FR, accounting for 26.38%, while T2T_frag has the lowest, representing only 1.43% (Table 2). Among the five T2T genomes, T2T_musa has the greatest number of FR on each chromosome (13,234), while T2T_homo has a comparatively small number (7,343). The T2T_mus, T2T_arab, and T2T_frag have even fewer, 443, 122, and 270, respectively (Table S3). These results indicate varying degrees of newly discovered sequences in the T2T genomes of different species.

The distribution pattern of FR is observed consistently in each T2T genome, showcasing similarities across different species (Figure 1 and Figure S1). In the case of the T2T_homo genome, extensive FR are detected in the centromere and telomere regions of the two chromosomes. Previous studies have indicated that in the T2T_homo genome, the centromere region in chr. 1 is located at 47% -59% of its total chromosome length, while the acrocentric region of chr. 15 is situated within the range of 0 - 23% of its chromosome position [26]. Remarkably, 64.6% and 69.2% of the centromere regions in chr. 1 and chr. 15, respectively, were identified as FR. Chr. 15, being one of the five acrocentric chromosomes in the human genome, was not sequenced before the release of the T2T genome. The FR are predominantly located in the acrocentric of chromosomes, accounting for 64.7% of the centromeres on average (Table S4 and Figure 1). For T2T_arab genome of *A. thaliana*, the FR are also extensively present at the centromeres and telomeres of the chromosomes (Table S5 and Figure S2B). The centromeres are located at approximately 45.6% to 53.0% of chr. 1 and 40.0% to 49.4% of chr. 5 [33]. Chr. 1 and chr. 5 of T2T_arab exhibit approximately 92.6% (chr.1) and 86.5% (chr.5) FR in the centromere regions, respectively.

**Figure 1.**
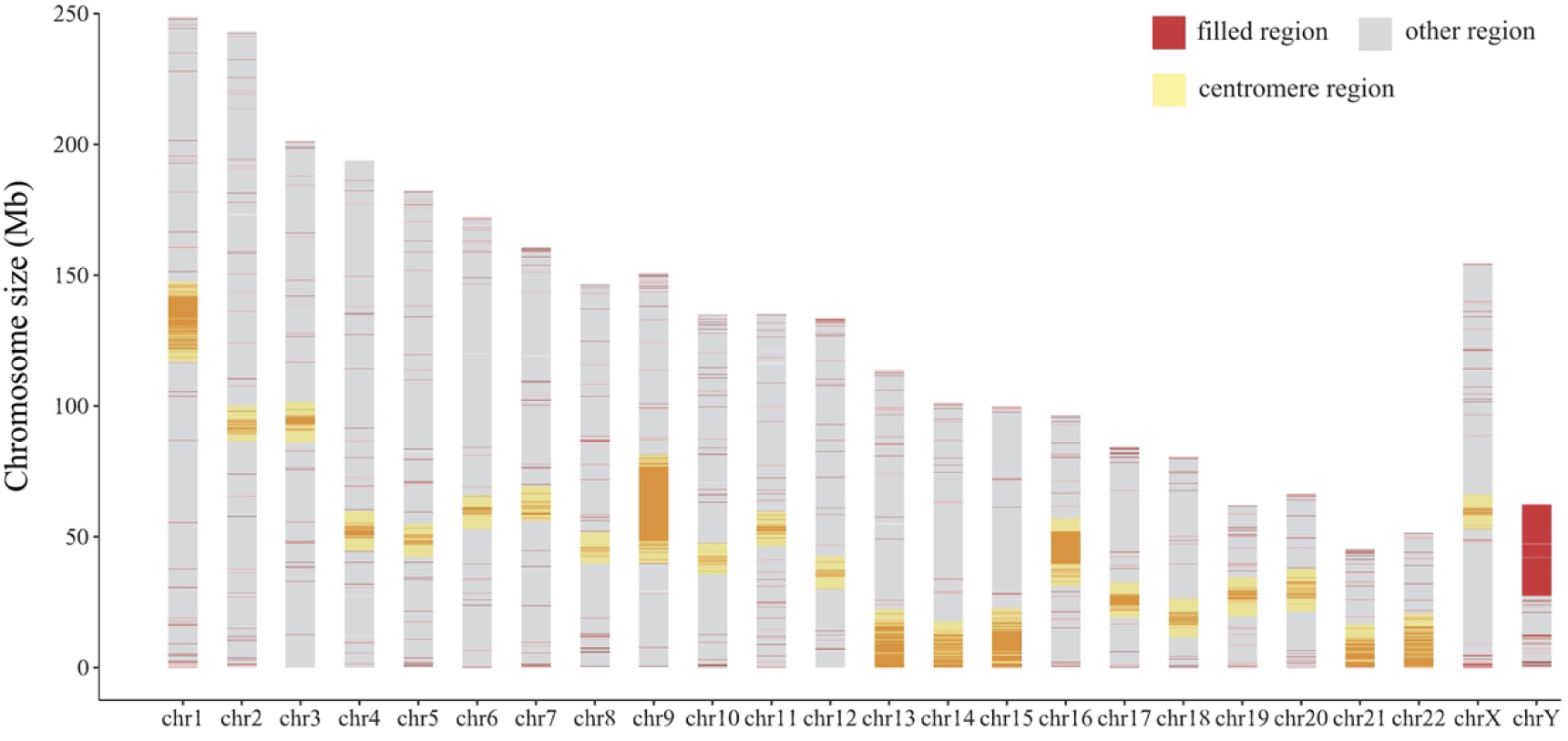
Distribution of FR in T2T genome of *H. sapiens*

On the other hand, the FR identified on chr. 4 of the T2T_mus genome are mainly distributed around the telomeres, with a higher concentration and density of FR loci near the telomere of the chromosome’s left end. The positions of FR loci on other chromosomes of T2T_mus suggest that the T2T genome primarily corrects and improves the FR mainly located around the telomeres of each chromosome, significantly enhancing the integrity and continuity of the *M. musculus* genome (Figure S2A). For chr. 3 of the T2T_musa genome of *M. acuminata*, a significant portion of FR is concentrated in the central region of the chromosome, coinciding with the site of centromere attachment. Additionally, there are numerous FR loci extending from the centromere towards both ends of the chromosome. Overall, the distribution of the FR on the chromosome gradually transitions from dense to sparse from the center to the sides. Furthermore, FR also appear at the telomeres of the chromosome. Compared to T2T_homo and T2T_arab, T2T_musa also has FR around the centromeres, which is attributed to the substantial improvement in the continuity of its genomic components achieved by Musa_pahang_v4 (T2T_musa) on the foundation of the ASM31385v2 (Old_musa) genome [31] (Figure S2C). The filling of numerous gaps in the assembly of the old genome is evident, as confirmed by the increase in N50 from 1.3Mb to 33.6Mb (Table 2). The continuity and integrity of the T2T_musa genome were significantly improved compared to the old genome.

The length of chr. 4 of F._vesca_v6 (T2T_frag) genome is 34,388,015 bp, and the left and right telomeres of chr. 4 are located at 1-1,078 bp and 34,387,198 - 34,388,015 bp, respectively [35]. And 19.0% and 6.9% of FR were identified in these two regions, respectively. The centromere sites were in 44.0% to 44.2% of the chromosome, and 95.5% of the centromere regions were newly filled. Meanwhile, partially FR are distributed in the remaining sites at the centromere and telomere. Observing at the distribution sites of the covered regions on the other chromosomes of the T2T_frag genome, the FR appeared in the centromere or telomere regions of each chromosome. Specifically, the telomere regions of chr. 1, 2, 5, and 7 were all identified as FR. (Table S6 and Figure S2D). Overall, the comprehensive analysis of FR suggests that the distribution of FR in different species may be associated with chromosome structure and function. A high occurrence of FR at the loci of centromeres and telomeres were consistently observed in human, animal and plant species.

To further unravel the genomic feature of FR, these regions were categorized into gene regions and intergenic regions for analysis. The proportion of intergenic regions was significantly higher than that of gene regions (exon and intron). In *H. sapiens*, the intergenic region accounts for 86.34%, while exon, CDS and intron account for only 0.23%, 1.99% and 13.26%, respectively. This trend is consistent across the remaining four species. For *H. sapiens*, *M. musculus, M. acuminata and F. vesca,* the proportion of intron in the FR is higher than that of CDS and exons (Table S7). In terms of specific types, the largest number of FR were identified as intron in *H. sapiens*, exon in *M. musculus* and *A. thaliana*, and intergenic in *M. acuminata*. Overall, the difference about the number of the four types in the FR of the three plants (*A. thaliana*, *M. acuminata* and *F. vesca*) was relatively small. However, there was a notable difference in the number between *H. sapiens* and *M. musculus* (Figure 2 and Table S7).

**Figure 2.**
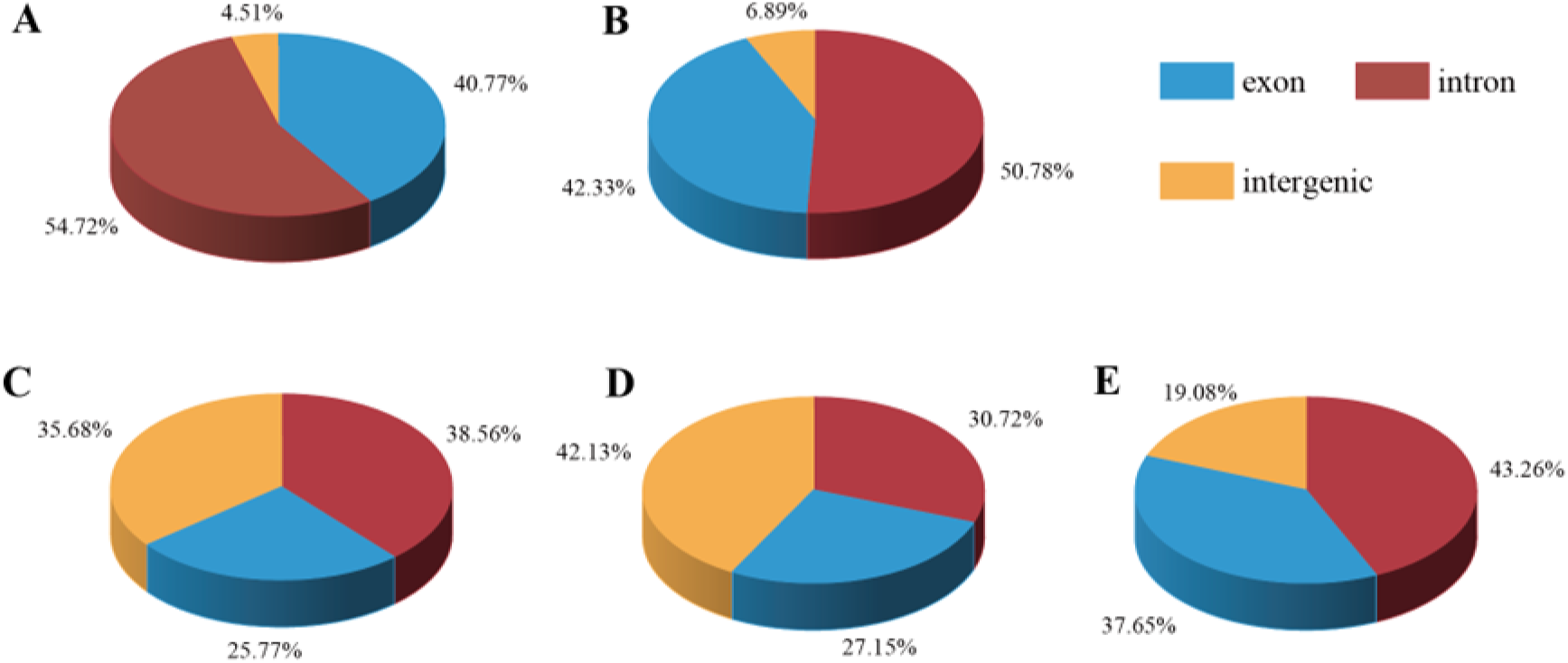
FR in genomics features (exon, intron, and intergenic). (A: *H. sapiens*; B: *M. musculu*s; C: *A. thaliana*; D: *M. acuminata*; E: *F. vesca*)

### 3.3 Analysis of the features of filled sequences

We annotated five T2T genomes for repeat sequence and conducted a detailed analysis of the features of repeat sequence within the FR. Repeat sequence includes DNA transposons, LTRs, LINEs, SINEs, and four other repetitive types. The proportions of repetitive sequences in the T2T genomes vary among different species. Results indicate that T2T_homo (*H. sapiens*), T2T_musa (*M. acuminata*) and T2T_mus (*M. musculus*) have relatively high proportions of repeat sequences, accounting for 54.60%, 52.37%, and 43.05%, respectively. While T2T_frag (*F. vesca*) and T2T_arab (*A. thaliana*) have a lower proportion of repetitive sequences, accounting for 35.56% and 24.00%. Additionally, there are differences in the proportions of different types of repetitive sequences within each T2T genome. In the T2T_homo genome, LINEs and SINEs are the predominant repetitive types. In the T2T_mus genome, LINEs and LTRs are the major repetitive types, with a significant proportion of SINEs as well. T2T_arab is mainly characterized by Satellite repetitive sequences. For both T2T_musa and T2T_frag, LTRs are the most frequent repeat type, accounting for 33.2% and 9.12%, respectively, while the proportions of the other types are relatively low. There were 14.85% and 18.98% of the repetitive types in T2T_musa and T2T_frag genomes, respectively, were unclassified. (Table 4 and Figure S3).

**Table 4.** DNA repeat sequences of T2T genomes in five species.

| Types | Genome proportion (%) |  |  |  |  |
| --- | --- | --- | --- | --- | --- |
|  | T2T_homo | T2T_mus | T2T_arab | T2T_musa | T2T_frag |
| DNA transposons | 3.54 | 1.05 | 4.00 | 0.60 | 3.32 |
| LTR | 8.64 | 11.64 | 6.35 | 33.20 | 9.12 |
| LINEs | 20.15 | 19.56 | 0.90 | 1.43 | 1.60 |
| SINEs | 12.60 | 7.26 | 0.06 | 0.56 | 0.50 |
| Simple_repeat | 3.14 | 2.61 | 1.09 | 1.32 | 1.12 |
| Satellite | 5.21 | 0.16 | 9.41 | 0.03 | 0.05 |
| Small RNA | 0.94 | 0.06 | 0.18 | 0.23 | 0.65 |
| Low_complexity | 0.20 | 0.37 | 0.33 | 0.15 | 0.22 |
| Unclassified | 0.18 | 0.34 | 1.68 | 14.85 | 18.98 |
| All | 54.60 | 43.05 | 24.00 | 52.37 | 35.56 |

Meanwhile, we observed correlation and differences in repetitive sequence distribution between the five species, indicating that DNA transposons have a higher proportion in T2T_homo, T2T_arab and T2T_frag LTRs are significantly more abundant in T2T_musa compared to other genomes. LINEs, SINEs and Simple repeats are highly expressed in T2T_homo and T2T_mus, while their proportions are lower in the remaining two plant species. Satellite has the highest proportion in T2T_arab, followed by T2T_homo. Small RNA has a higher proportion in T2T_homo and T2T_frag compared to the other four genomes. Low complexity has a higher proportion in T2T_mus and T2T_arab compared to other genomes. Overall, the proportions of different repetitive sequence types are relatively similar in T2T_homo and T2T_mus genome, while T2T_arab, T2T_musa and T2T_frag show similar patterns in terms of repetitive sequence composition (Figure 3A).

**Figure 3.**
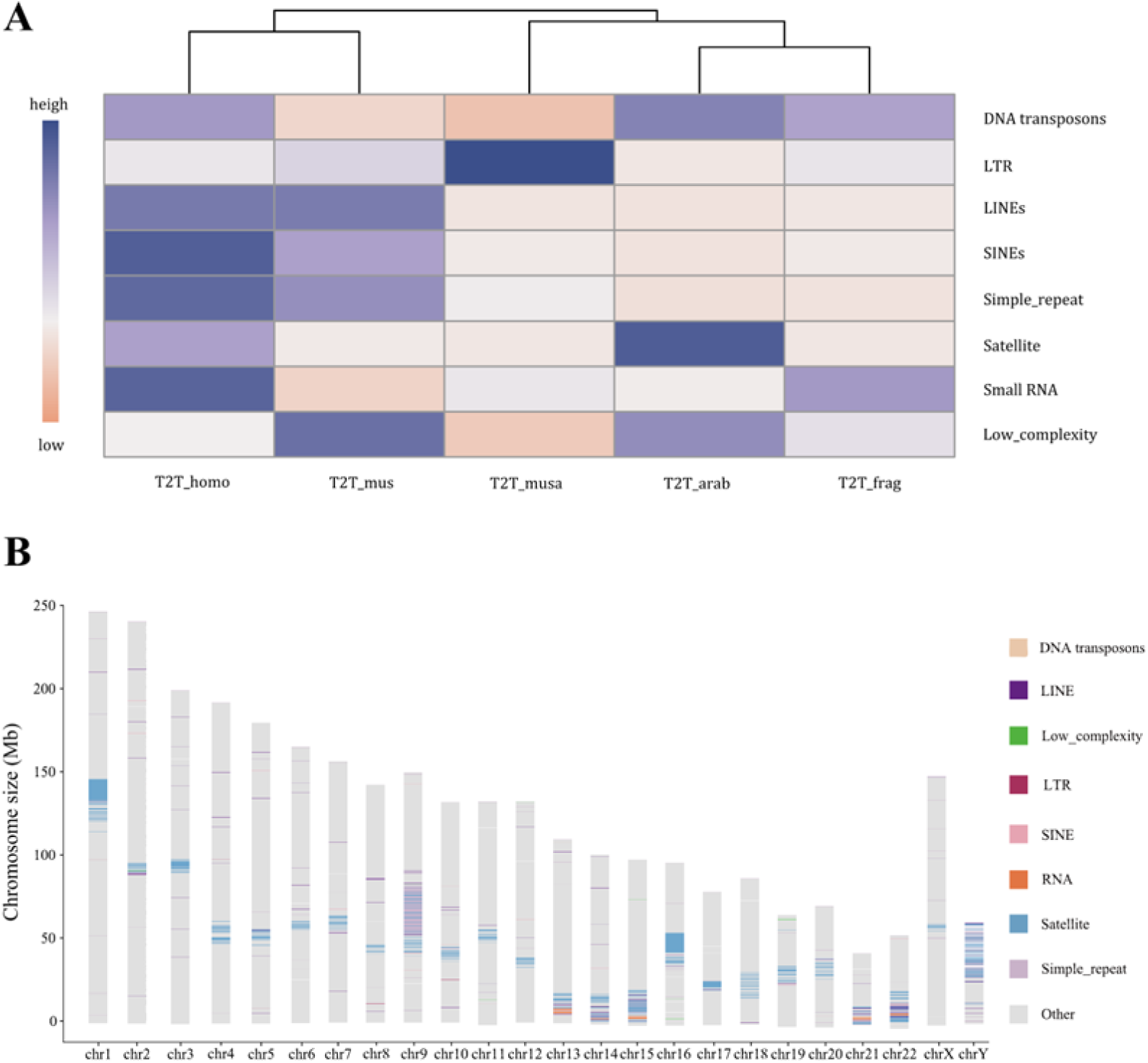
Repetitive Sequences in five T2T genomes. (A: Clustering of repeat types in T2T Genomes; B: Distribution of repeat types within FR of *H. sapiens*)

Based on the identification of FR and the annotation of repetitive sequences, we extracted the repetitive sequence that appeared in all FR. The FR of T2T genomes contained a substantial number of repetitive sequences. In T2T_homo, the FR of chr. 1 and chr. 5 contains 93.14% and 86.96% of repetitive sequences, respectively. Similarly, in FR of chromosomes represented by T2T_arab and T2T_musa, the proportion of repeat sequences exceeded 90%. In contrast, the proportion of repetitive sequences in FR of T2T_frag and T2T_mus is relatively low, accounting for 71.43% and 46.31%, respectively (Table S8). In the FR of chr. 1 and chr. 15 in T2T_homo genome, they are primarily covered by Satellite and Simple repeat, constituting 82.8% and 6.26% in chr.1 and 45.23% and 31.49% in chr. 15, respectively. There are also a few LINEs and SINEs elements covering their FR. For chr. 4 in T2T_mus genome, we observed that the upper part of its FR contains a higher density of repetitive sequences, mainly composed of LINEs (15.7%) and LTR (10.71%). In chr. 1 and chr. 5 of T2T_arab genome, a large number of Satellite sequences cover the majority of their FR, accounting for 92.14% and 83.05%. Examining chr.3 in T2T_musa and chr. 4 in T2T_frag, we found that LTRs, account for the majority of their FR loci (55.64% for T2T_musa and 36.57% for T2T_frag), with relatively many SINEs in the FR of these two chromosomes (9.02% for T2T_musa and 1.45% for T2T_frag) (Table S9, Figure 3B and Figure S3).

### 3.4 Gene annotation of FR

The gene information of the five T2T genomes, both on the entire genome and within the FR, was identified to analyze their gene function. The results indicated significant differences in the number of genes present in the entire genome and FR among five T2T genomes. T2T_homo and T2T_mus have a considerably higher number of identified genes in the entire genome compared to the other three genomes, while the number of genes identified in T2T_arab is lower than the other genomes. Regarding the number of genes in FR of the genomes of five species, T2T_homo and T2T_musa have a higher number of genes than the other 43genomes, with 5,930 and 5,157 in their FR, respectively. On the other hand, T2T_arab has a considerably lower number, with only 251 genes. The proportion of genes in the FR relative to the entire genome varies among five T2T genomes. T2T_homo and T2T_musa genome have a relatively high number of genes in the FR compared to the entire genome, accounting for 14.3% and 14% of the entire genome, respectively. T2T_mus, T2T_frag and T2T_arab have relatively fewer genes in their FR, accounting for only 2.3%, 1.34% and 0.75% of the entire genome, respectively (Table S10).

Among the annotated genes in T2T genome, different numbers of unique annotated genes were found in *H. sapiens*, *M. musculus* and *A. thaliana*, with 5,827 unique annotated genes in mouse and only 33 newly annotated genes in *A. thaliana*. 75.82% of the unique annotated gene regions in *H. sapiens* were identified as repeats, while only 36.63% in *M. musculus* and 2.01% in *A. thaliana* were repeats (Table 5 and Table S11). The different metabolic pathways for the newly annotated genes were analyzed in *M. musculus*. The results of KEGG enrichment analysis showed that some genes were found in the pathways of histamine metabolism, sulfur metabolism, RNA polymerase, tryptophan metabolism, Systemic lupus erythematosus (SLE), arginine and proline metabolism, circadian rhythm, etc. Among them, the number of genes enriched in SLE was the highest (Figure S4).

**Table 5.**
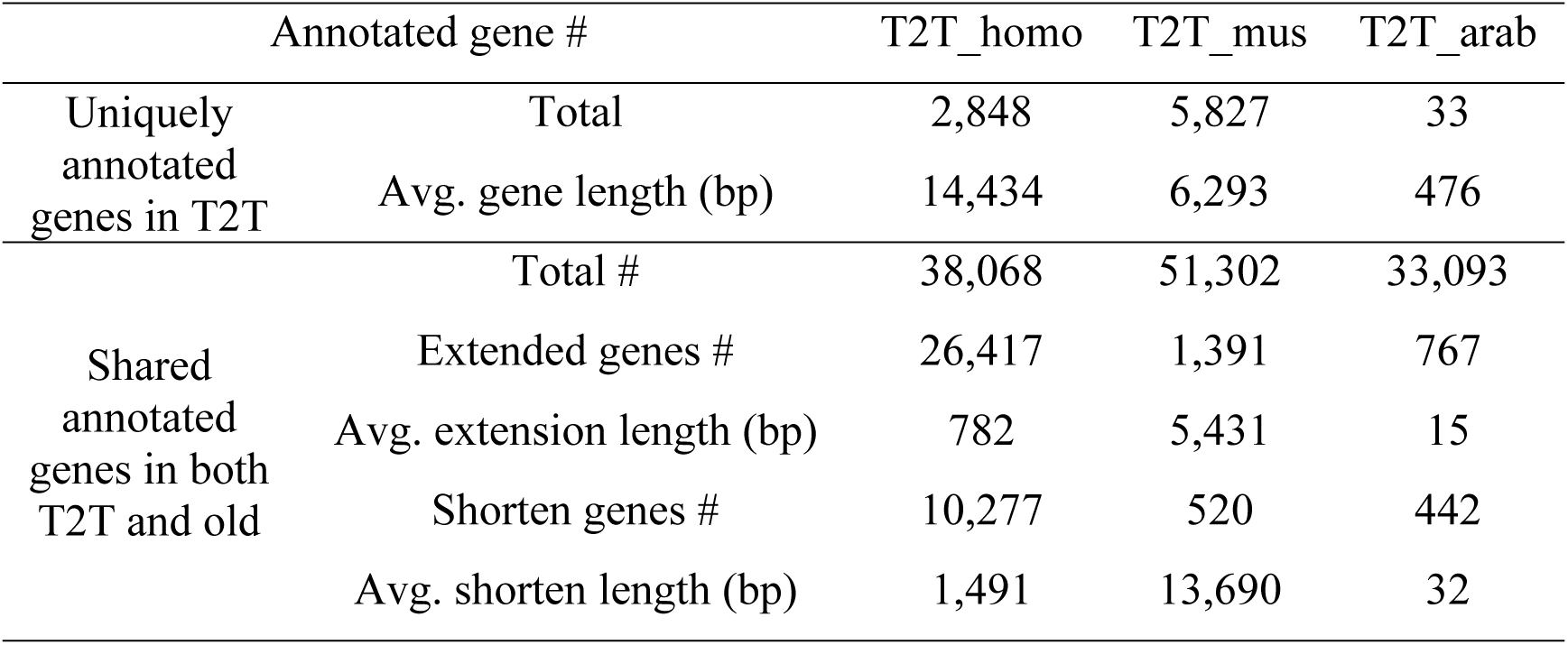
Statistics of gene annotation in T2T genome.

For the common annotated genes of the T2T genome and the old genome, the length of each pair of identical genes is not fixed. There are 26,417, 139 and 767 common annotated genes have been extended in the T2T_homo, T2T_mus and T2T_arab, respectively, while others have been shortened or remained unchanged. In *H. sapiens*, *M. musculus* and *A. thaliana*, the number of extended genes was greater than that of shortened genes, and the number of extended genes was 2.6, 2.7 and 1.7 times that of shortened genes, respectively. The average number of extended gene bases is smaller than the shortened gene bases. The average length of shortened bases in *H. sapiens*, *M. musculus* and *A. thaliana* is 1.9, 2.5 and 2.1 times that of extended bases, respectively. The *GPAT2* gene exhibits the most significant extension in the T2T genome of *H. sapiens*, stretching by 608,848 bp compared to the old genome. It is located on chr. 2 from 95,919,724 bp to 96,542,891 bp. The *GPAT2* gene encodes glycerol-3-phosphate acyltransferase 2, a crucial enzyme involved in the triglyceride synthesis pathway during fat metabolism. *GPAT2* is a mitochondrial isoform that is mainly expressed in the testis under physiological conditions. It is aberrantly expressed in multiple myeloma and is identified as a novel cancer testicular gene [57]. The TMEM267 gene, located in chr. 13 of T2T_mus genome from 119,624,575 bp to 120,072,595 bp, has experienced the longest extension, with an increase of 325,000 bp compared to the old genome. This gene encodes a transmembrane protein (*TMEM267*) predicted to be a component of cell membrane. It shares homology with the human TMEM267 and is generally expressed in adult testis, the central nervous system and other tissues [58]. In the T2T genome of *A. thaliana*, the *AT2G43150* protein coding gene located between 20,458,156 bp and 20,462,774 bp on chr. 2 has the longest extension compared to the old genome. The gene ID 818917, Proline-rich extension-like family protein, acts in cell wall and participates in plant cell wall organization [59, 60].

For the RNA-seq data alignment results on the T2T genome, the average alignment ratio for each species exceeded 80%. As for the FR, *H. sapiens* and *F. vesca* exhibited the highest alignment number (4,407,560 and 8,738,360) and ratio (14.1% and 13.2%), respectively. *A. thaliana* demonstrated a relatively lower alignment number and ratio of 226,987 (1.9%) compared to its counterparts (Table 6).

**Table 6.** Statistics of mapping results between RNA-seq data and T2T genome.

|  | T2T_homo | T2T_mus | T2T_arab | T2T_musa | T2T_frag |
| --- | --- | --- | --- | --- | --- |
| SRA accession | DRP011239 | SRP492204 | SRP470572 | SRP102870 | SRP115444 |
| Total reads # | 277,824,945 | 90,365,441 | 130,090,066 | 74,536,889 | 292,401,740 |
| Avg. MP | 80.5% | 99.6% | 98.7% | 99.0% | 99.5% |
| Avg. MN of FR | 4,407,560 | 1,703,233 | 226,987 | 2,165,737 | 8,738,360 |
| Avg. MP of FR | 14.1% | 9.0% | 1.9% | 10.4% | 13.2% |

(Avg. MP: Average mapping ratio; Avg. MN of FR: Average mapping numbers of filled regions; Avg. MP of FR: Average mapping ratio of filled regions)

### 3.5 Gene function and reliable FR of *H. sapiens*

Building on the validation of the authenticity of FR identification and its potential transcriptional activity, we conducted a comprehensive functional prediction analysis of the FR sequences. This analysis identified 1,007 enriched motif signature sequences, among which 172 were particularly significant. Of these, 151 motifs exhibited p-values and q-values approaching 0, indicating strong statistical significance. Notably, the sequence "AAATATCT" was associated with multiple motif signatures, including "EPR1 (MYB-related)/colamp-EPR1-DAP-Seq," "LHY1 (MYB-related)/col-LHY1-DAP-Seq," "AT3G10113-DAP-Seq," and "At4g01280 (MYB-related)/colamp-At4g01280-DAP-Seq". In addition, the sequences "KTGTTTGC" and "AAGATTCT" were found to be highly enriched, comprising 67.87% and 50.46% of the target sequences, respectively, compared to 48.76% and 32.91% in the background sequences. Existing research suggests that these motif signature sequences may play critical roles in regulating gene expression or participating in specific regulatory pathways. They are also associated with species-specific functional pathways and identification, further highlighting the importance of FR sequence identification (Table S12).

To explore the functional relevance of the 172 significant motif sequences, we performed KEGG pathway enrichment analysis, which revealed 45 enriched pathways. Among these, eight pathways, including the "Hippo Signaling Pathway -Multiple Species," "Cellular Senescence," and "Inflammatory Bowel Disease", demonstrated significant enrichment (Figure 4).

**Figure 4.**
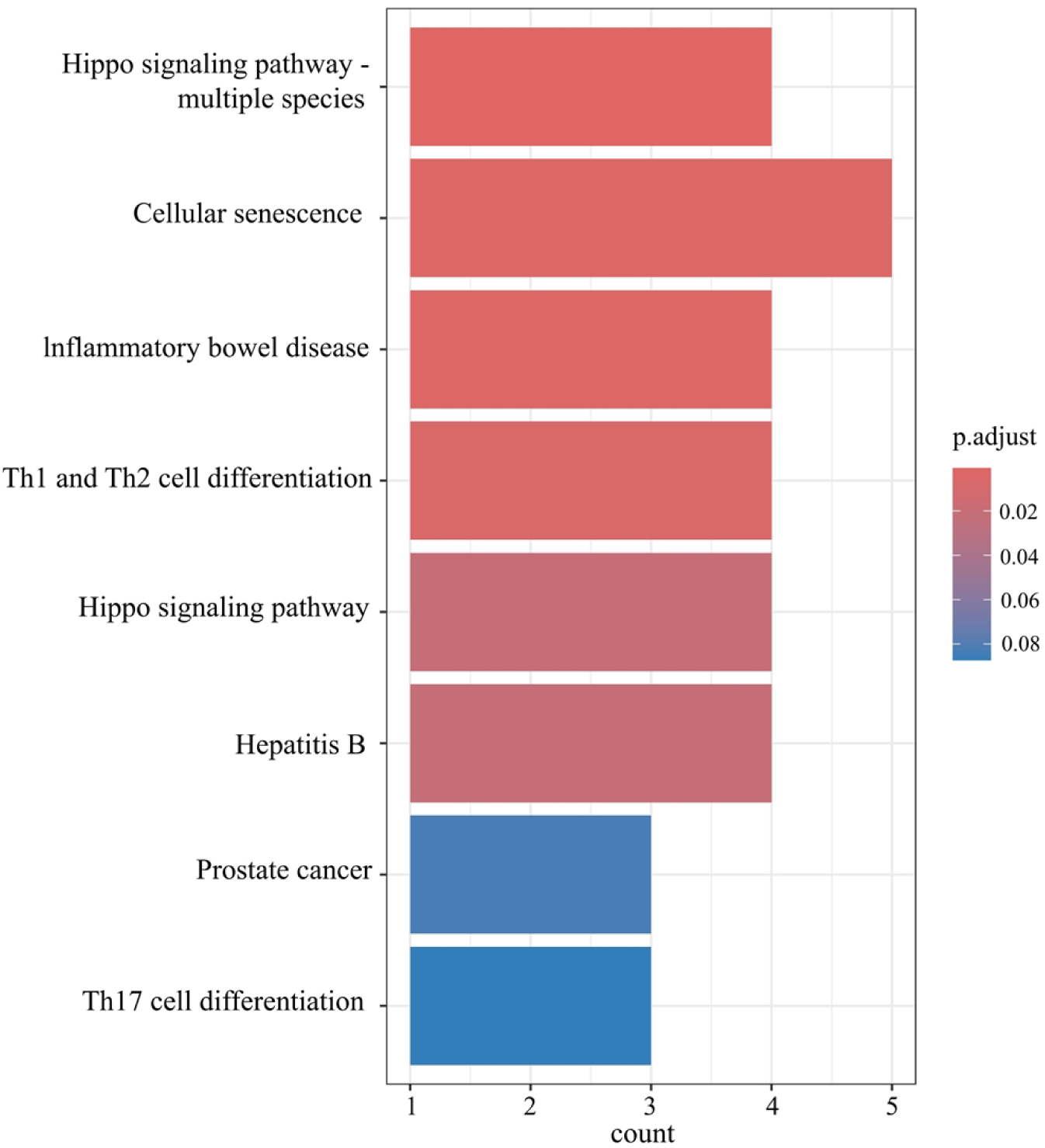
KEGG enriched pathway statistics for significant motif genes for *H. sapiens*

The Hippo signaling pathway primarily regulates cell proliferation, apoptosis, and tissue growth by modulating the activity of the transcription factors YAP and TAZ [61]. Disruption of this pathway is linked to the development of various cancers and plays a critical role in tissue development and repair. Inflammatory bowel disease (IBD), encompassing Crohn’s disease and ulcerative colitis, is a chronic inflammatory condition resulting from aberrant immune responses to intestinal microbiota or dietary antigens. The differentiation and balance of Th1 and Th2 cells are essential for optimal immune system function [62]. Th1 cells primarily combat intracellular pathogens, such as viruses and certain bacteria, through the production of interferon-gamma (IFN-γ) and other cytokines. In contrast, Th2 cells target extracellular pathogens and regulate allergic responses by secreting cytokines such as IL-4, IL-5, and IL-13.

The KEGG enrichment pathway analysis indicates that the genes associated with these motifs are implicated in processes related to cell proliferation, aging, immune response, and inflammation. These findings provide valuable insights for further research into the mechanisms underlying these biological processes and their associated diseases.

By comparing the RNA seq data successfully aligned to FR (denoted as FR_FILTER_RNA) with the NT database, we observed that the FR sequences mapped not only to the H. sapiens database but also to databases of Canis lupus familiaris, Leporidae, and Sus scrofa. Notably, there were significantly more sequences aligned to Leporidae and S. scrofa than to C. lupus familiaris, with 2,206 and 1,339 sequences, respectively (Table S13). This indicates that FR sequences exhibit homology and similarity across these species. We designated these sequences as FR_FILTER_RNA_OTHER-SPECIES and subsequently aligned them to the GRCh38 genome. During this alignment, seven sequences remained unaligned, which were deemed as the most reliable sequences (Table S14).

Among the seven candidate sequences, we ultimately selected sequences 4, 5, and 7 for bidirectional PCR amplification and Sanger sequencing. Sequences 3 to 7 are located in the centromeric regions of chromosomes 21 and 22, with sequences 4 and 5 exhibiting the highest number of mapped RNA-seq samples (both 13). These two sequences were selected to facilitate the comparison of sequence characteristics across different centromeric regions, while sequence 7, compared to sequence 6, was prioritized as it is closer to the centromeric core region and may exhibit higher transcriptional activity or important RNA-binding sites. The final selected sequences were NC_060945.1 (3652960–3654953, FR_seq_1), NC_060946.1 (5018082–5019176, FR_seq_2), and NC_060946.1 (5627132–5628767, FR_seq_3). Specific primers were designed for PCR amplification, yielding theoretical fragment lengths of 594 bp (FR_ seq_1), 990 bp (FR_ seq_2), and 1383 bp (FR_ seq_3). Electrophoresis results showed single, clear bands for all three sequences, with band positions fully consistent with the expected fragment lengths, indicating the absence of non-specific contamination. Notably, FR_ seq_2 and FR_ seq_3 exhibited clearer and brighter bands compared to FR_ seq_1 (Figure 5A). Following electrophoretic separation, fluorescent detection and data acquisition were performed, generating chromatograms for the three sequences. The chromatogram results demonstrated that the bases in the filled regions, the original sequences, and their junctions were accurate and free of mismatches. The peaks were sharp, well-separated, and evenly distributed, with base signal intensities uniform and peak colors and orders consistent with the expected sequences.

**Figure 5.**
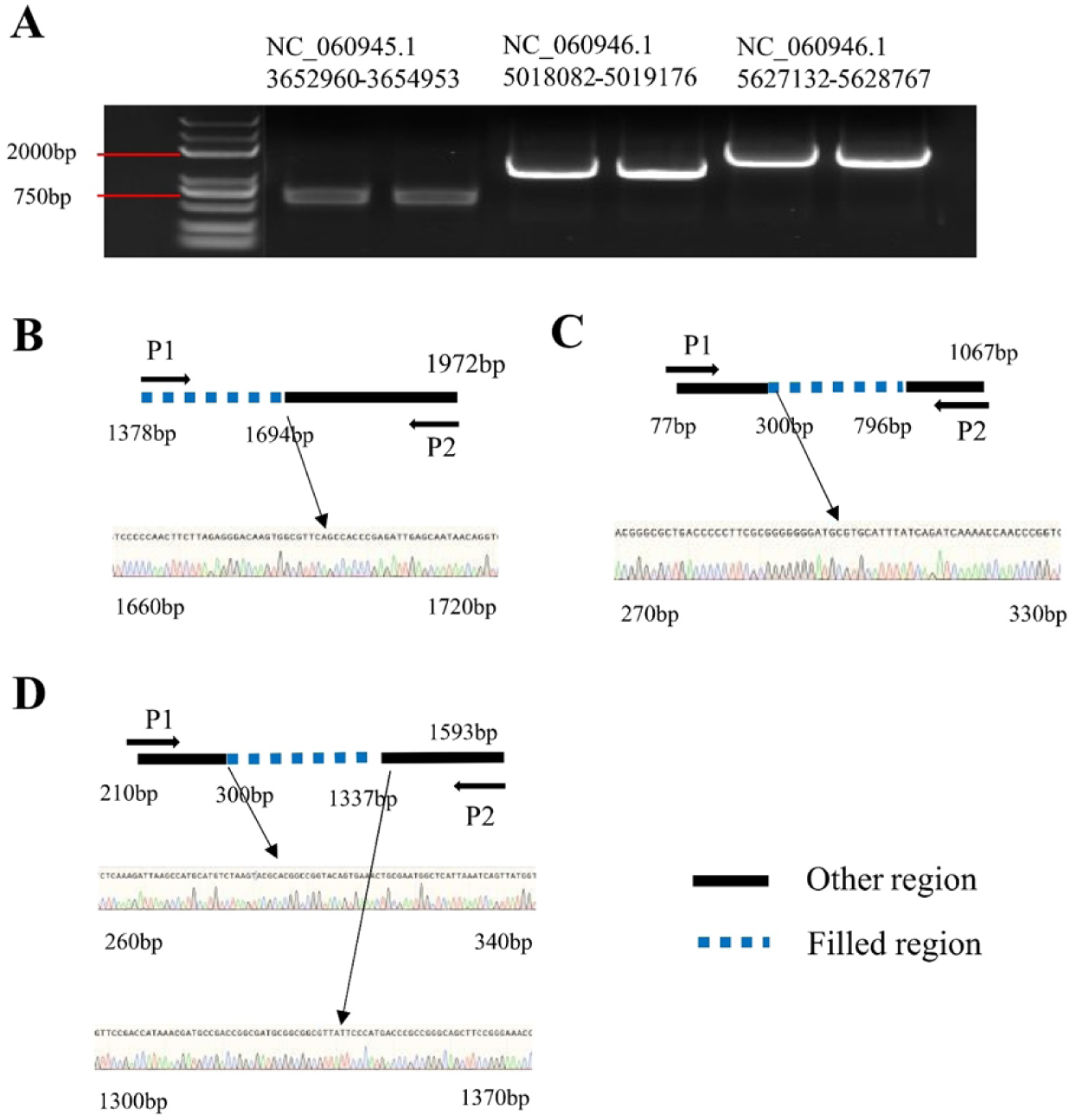
Sanger sequencing results of three reliable sequences (A: Electrophoresis analysis of PCR amplification products for three target sequences; B-D: Sanger sequencing assembly results for three target tequences) *(NC_060945.1 3652960-3654953 as FR_ seq_1, NC_060946.1 5018082-5019176 as FR_ seq_2, and NC_060946.1 5627132-5628767 as FR_ seq_3)*

Specifically, the junction regions for FR_ seq_1 (1660 bp–1720 bp), FR_ seq_2 (270 bp–330 bp), and FR_ seq_3 (260 bp–340 bp) were highly consistent and matched perfectly with the reference sequences. Collectively, these results confirm the authenticity of FR_ seq_1, FR_ seq_2, and FR_ seq_3, providing reliable data to support subsequent functional analyses (Figure 5B-5D).

## 4. Discussion

By conducting pairwise comparisons of the T2T and old genomes across five species, this study has systematically identified and analyzed the proportions, locations, properties, and gene functions of FR. This comprehensive analysis aims to unveil differences and commonalities in genome composition, structure, and functionality. Various T2T genomes have been successively published for human, animals and plants. Although previous studies have explored these features in specific organisms [26, 29, 31, 33, 35, 63], there are a cohesive examination comparing representative model organisms across species. Our study not only longitudinally compared the old and T2T genomes, but also horizontally compared the similarities and differences across T2T genomes from diverse species. Through horizontal comparison, it was concluded that the FR of T2T genome has distinct pattern characteristics in human, animal and plants (Table 4 and Figure3). This study offers valuable insights into shared characteristics among different species in the complete genome assembly. It provides a deeper exploration of biodiversity and the evolutionary process, contributing to a more comprehensive understanding of genomics.

The T2T genomes of the five species exhibit higher quality and completeness compared to their old genomes. They contain a greater number of newly discovered sequences and functionalities. This indicates significant advancements in sequencing and assembly technologies for the T2T genomes. These improvements enable more accurate representations of biological realities. Evolutionary processes have subjected different species to distinct selective pressures and genomic variations, resulting in variations in chromosome structures [64, 65]. In specific comparisons, such as *H. sapiens*, *M. musculus* and *F. vesca*, changes in genome size and GC content are relatively minor when comparing T2T genomes to the old genomes. However, significant changes are observed in *M. acuminata* and *A. thaliana* when compared to their old genomes (Table 2). These differences underscore the species-specific nature of genomic evolution and highlight the importance of advanced technologies in capturing these variations comprehensively.

The analysis of location distribution and the sequence characteristic of FR reveals consistent patterns across these five species, which highlights the intricate connection between the distribution of FR and chromosome structure and functionality. In the T2T genomes of *H. sapiens* and *A. thaliana*, the FR are predominantly located in telomeres and centromeres, underscoring the vital role of these structures in genome variation and evolution, essential for maintaining genome stability and integrity (Table S4-S6 and FigureS2). A notable finding is the prevalence of repetitive sequences within these FR, particularly evident in the T2T genomes of *H. sapiens*, *A. thaliana* and *M. acuminata* (Table S8 and FigureS3). It indicates that the FR may result from insertions or amplifications of repetitive sequence, indicating a potential contribution of repetitive sequences to innovation in genome structure or functionality. Sergey Nurk et al demonstrated that most of the unsequenced parts of the short arms of the five acrocentric chromosomes of *H. sapiens* were enriched by satellite repeats and segmental duplications [26]. Matthew Naish et al fully assembled the centromeres of *A. thaliana* and identified millions of CEN180 satellite base pairs in the centromeric region, supporting CENH3 loading [33]. Our analysis aligns with these conclusions, revealing similar patterns in multiple model organisms.

In gene annotation analysis of the FR, the number of genes present in the FR and the whole genome was quantitatively compared. Delving into the intricacies, we explored the divergence in annotation information between the T2T and old genomes, thereby identifiying newly annotated genes in the T2T genomes. In the case of common annotated genes, we also examined the performance in the two genome versions. *H. sapiens* and *M. musculus* exhibited a higher number of genes in the whole genome, while *A. thaliana* displayed a comparatively smaller count (Table S10). This observation indicates disparities in gene functions and complexity among species, both in the whole genome and the FR. The newly annotated genes in *M. musculus* were notably enriched in pathways related to systemic lupus erythematosus and other functions, indicating that the newly annotated genes may be related to biological performance metabolism and diseases. For the co-annotated genes, GPAT2 [57], TMEM267 [58] and AT2G43150 [59, 60] genes were with the largest length variation among *H. sapiens*, *M. musculus* and *A. thaliana*, respectively (Table 5). These genes likely play pivotal roles in maintaining genome integrity and functionality. For *M. acuminata* and *F. vesca* genomes, due to the different gene symbols in annotation files of the T2T genome and the old genome, coupled with the customization of all gene annotation information, we refrained from conducting detailed statistical analyses on the annotated gene changes in these species. We aligned RNA-seq data to the T2T genome (Table 6). By mapping RNA-seq data to the FR of the T2T genome, it further confirms the possible presence of transcription in the FR and illustrates the necessity of in-depth investigation of the filled region of T2T genomes.

The T2T genome of human (T2T-CHM13v2.0) assembly, compared to the old (GRCh38), has 60.7% of FR located on the Y chromosome (Table S3 and Figure1). It is significantly higher than that of other chromosomes. Due to the complex structure and repetitive regions, a significant portion of the Y chromosome remained incompletely assembled [26, 27]. The initial release of the complete human genome by T2T involved the assembled the 22 pairs of autosomes and the X chromosome in their entirety. Subsequent efforts by the T2T consortium successfully addressed this challenge by assembling all chromosomes from a complete hydatidiform mole genome. This comprehensive assembly not only resolved over 50% of the base gaps in the Y chromosome of the GRCh38 genome and but also corrected multiple errors in GRCh38-Y [66]. In addition, the proportion of FR on the chr. Y in *M. musculus*, accounting for 21.4%, further underscores the advancements in sequencing technologies for T2T genomes in different species. These advancements signify a gradual overcoming of challenges in genome assembly, paving the way for the completion of previously gap genomes.

We conducted an in-depth comparison of the sequencing and assembly methods between the T2T and prior genome versions for five species. Initially, we collected the assembly strains or samples corresponding to each genome version. Both the T2T and the old genome assembly of *H. sapiens* used the ’complete hydatidiform mole’ sample. Specifically, the T2T genome used the CHM13 sample, while the GRCh38 genome combined CHM1 and CHM13 samples [26, 27]. For *M. musculus*, *A. thaliana*, *M. acuminata*, and *F. vesca*, the different versions were sequenced and assembled from the same samples or strains. For instance, the GRCm39 and GRC38 versions of the *M. musculus* genome both used the ’C57BL/6J’ strain [28, 29]. Similarly, the two versions of *A. thaliana*, *M. acuminata*, and *F. vesca* genomes used the Col-0 (Columbia) ecotype (32, 33), Double haploid Musa acuminata spp. malaccensis (DH-Pahang) [30, 31], and ’Hawaii 4’ [34, 35], respectively (Table S15).

We also examined the sequencing and assembly methods of the two genome versions. The T2T genome’s methods are more comprehensive and advanced, leading to the identification of different numbers and proportions of FR in the T2T genome. For the T2T genomes of all five species, both short-read and long-read sequencing methods were used. The prior *H. sapiens* genome used Sanger sequencing, Illumina short-read sequencing, Pacific Biosciences (PacBio) long-read sequencing, and Oxford Nanopore Technologies (ONT) nanopore sequencing [27]. In addition to these methods, the T2T genome also used BioNano optical maps and single-cell DNA template strand methods [26]. The sequencing methods for the two versions of *A. thaliana* were similar, but different assembly tools were used: the old version employed tools like Trimmomatic and SPAdes, while the T2T version used Guppy, Porechop, Filtlong, and Flye for base calling, data processing, and genome assembly [32, 33]. For *M. acuminata* and *F. vesca*, the old version used 454 and Illumina sequencing methods, whereas the T2T incorporated long-read sequencing (ONT sequencing) [34, 35] (Table S16). Despite the differences in sequencing and assembly methods between the T2T and old genome, they were derived from the same assembly samples. This commonality ensures a consistent reference point for comparing different versions.

During the comparison, we used cross-validation and consistency analysis tools such as MUMmer [42], minimap2 [53], and BUSCO [24]. We also integrated highly similar functional annotations and homology analyses of different versions to ensure the accuracy of our FR identification results. For instance, considering the two versions of the human genome, we aligned them using both MUMmer and minimap2, and then extracted the overlapping regions in the FRs generated by each software. By calculating, we found that the average overlap rate of FR_2 (FR obtained by minimap2) in FR_1 (FR obtained by MUMmer) reached 78.6% (Table S17). This confirmed that, despite using different alignment software, the FR obtained were highly similar, further verifying the authenticity and reliability of these FRs. Meanwhile, the enhanced integrity of the T2T genome can also be attributed to the advancement of sequencing and transcriptomic tools.

To further substantiate this conclusion, we combined it with previous RNA-seq alignment results on the T2T genome. We extracted the regions where RNA-seq aligned in both FR_1 and FR_2 and conducted quantitative analysis. We discovered that among the 5535 FRs identified in the T2T genome, 3582 FR regions were successfully aligned by RNA-seq. For instance, in chromosome 5, a total of 233 FR sequences were detected, with 176 successfully aligned by RNA-seq (Table S18). Additionally, we counted the coverage of RNA-seq after alignment in the FR. The results indicated that in chromosome 9 and chromosome 1, 66.9% and 68.7% were mapped to the FR, with coverage rates of 91.2% and 82.6%, respectively. On average, RNA-seq coverage of FRs across all chromosomes was 53.5% (Table S18). These results strongly support the correctness of FR identification and additionally confirm potential transcripts and transcriptional activity in the FRs. For FRs mapped by a large number of reads, there may be higher gene expression abundance and levels.

Compared to the old version of reference genomes, the T2T genomes represent a significant advancement. This comprehensive study systematically explored the genomic features within FR of reference genomes across five model species. A nuanced examination of mutations and polymorphisms at the individual level emerges as pivotal for deciphering genetic variations, pinpointing diseases, and refining strategies for animal and plant breeding. Future endeavors should aim for in-depth comparisons within individual genomes, contextualized within the broader landscape of the pangenome, to unveil more intricate patterns and insights.

## Author contribution statement

ZL conceived and designed the study and supervised the data analysis. CX conducted data collection, software execution, data analysis, and manuscript writing. JZ, LHH and ZL revised the manuscript and guided the formatting of papers. CX, HZ, YPZ, MW proofread the paper format.

## Data availability

The T2T and old genome assemblies of *H. sapiens*, alongside the old genomes of *A. thaliana* and *M. acuminata*, are accessible via NCBI’s RefSeq repository (https://www.ncbi.nlm.nih.gov). The RefSeq accession numbers for these assemblies are as follows: GCF_000001405.40 and GCF_009914755.1 for the old and T2T genome assembly of *H. sapiens*, GCF_000001733.4 and GCF_000313855.2 for the old genome of *M. acuminata* and *A. thaliana*. Comprehensive data on the remaining T2T and old genome assemblies can be acquired by consulting the details provided in Table 1. The RNA seq dataset information of the five species can be obtained through the SRA accession of DRP011239, SRP492204, SRP470572, SRP102870 and SRP115444, respectively. The specific BioProject and sample information of RNA seq can be obtained through Table S1.

